# The DNA Mismatch repair protein, MSH6 is a novel regulator of PD-L1 expression

**DOI:** 10.1101/2024.11.28.625929

**Authors:** Kirsten Brooksbank, Charlotte Smith, Eleni Maniati, Amy Gibson, Wai Yiu Tse, Amy Kate Hall, Jun Wang, Tyson V Sharp, Sarah A Martin

## Abstract

Immune checkpoint inhibitors (ICIs) are extremely effective in a subgroup of mismatch repair-deficient (MMRd) cancers, but ∼50% remain resistant to treatment. We have shown for the first time that this may be due to the differential regulation of factors linked to response to ICIs upon loss of the different MMR genes. Here, we show that increased PD-L1 expression is observed upon loss of the MMR genes MLH1, MSH2 and PMS2. However, this is not true upon loss of MSH6. Here, we show that this is due to a novel role for MSH6 as a direct regulator of PD-L1 transcription, dependent on recruitment by the histone trimethyltransferase SETD2. Next-generation sequencing of MLH1 and MSH6 knockout (KO) cells revealed that MSH6 KO cells have significantly lower microsatellite instability in comparison to MLH1 KO cells, despite MSH6 KO cells having a higher mutational burden. These findings emphasise the need for gene-specific stratification in the MMRd cohort.

## INTRODUCTION

The DNA mismatch repair (MMR) pathway is primarily concerned with the repair of DNA lesions that occur during DNA replication, such as base-base or insertion/deletion (indel) mismatches (Hsieh and Yamane, 2008). In humans, the MMR pathway is initiated by one of two different heterodimeric complexes of MutS-related proteins, MSH2-MSH6 (MutSα) or MSH2-MSH3 (MutSβ), each with a differing lesion recognition specificity. This interaction enables MutS to recruit the MutL complex, itself a heterodimer consisting of complexes of MLH1/PMS2 (MutLα) or MLH1/PMS1 (MutLβ), for the excision and repair of the damage. MMR-deficient (MMRd) tumours exhibit a mutator phenotype and are clinically characterized by high levels of microsatellite instability (MSI) (Hsieh and Yamane, 2008). This is a type of DNA damage that results in extensions and contractions in regions of the genome called microsatellites, which consist of short, repeated sections of nucleotides.

The use of immune checkpoint inhibitors (ICIs) has transformed the treatment of various solid tumours by blocking immunosuppressive signals to promote anti-tumour immunity. However, response rates are extremely variable necessitating the identification of predictive biomarkers for response. The most successful predictive biomarker for ICI response to date is MMRd/MSI-high (MSI-h), which received full approval for the treatment of metastatic or unresectable MSI-h/MMRd solid tumours with pembrolizumab in 2023 (Geoerger *et al*., 2020; O’Malley *et al*., 2022; Le *et al*., 2023). The high tumour mutational burden (TMB) in MMRd tumours is often credited with eliciting a strong immune response and, therefore, susceptibility to ICI treatment. However, resistance to anti-PD-1 therapy is observed in 40-70% of MMRd patients (Mestrallet *et al*., 2023), highlighting the need to further refine our understanding of the mechanisms driving response and resistance to ICIs to better identify precise predictive biomarkers. There is accumulating evidence that MMRd/MSI-h is too broad a biomarker and that this could be why such variable response rates are observed in the MMRd/MSI-h cohort. For example, an *in vivo* study observed that MSH2 KO tumours with low MSI had a worse response to anti-PD-1 therapy than matched MSH2 KO tumours with high MSI (Mandal *et al*., 2019). In addition, genomic analysis from ICI-treated MMRd CRC and esophagogastric patient tumours also revealed that heterogeneity in MSI levels in the MMRd population may contribute to the variable response rates to ICIs observed. This is consistent with other studies that have shown that MMRd tumours are not necessarily MSI-h, even though MMRd/MSI-h received joint approval as a biomarker for response to pembrolizumab (Vanderwalde *et al*., 2018).

Another important factor associated with response to ICIs is expression of the immune checkpoint protein Programme Death Ligand 1 (PD-L1). Pembrolizumab targets the PD-1/PD-L1 interaction and PD-L1 expression is utilised clinically to identify patients that may benefit from ICIs across several cancer types. PD-L1 expression is induced by both DNA damage and immune signalling, such as interferons (IFNs) and interleukins, via the JAK/STAT signalling axis (Mogensen, 2019; Sato, Jeggo and Shibata, 2019). Therefore, there is a logical link between MMRd/MSI-h and elevated TMB leading to increased PD-L1 expression, but there is conflicting evidence on whether this is the case. For example, numerous studies have demonstrated that PD-L1 expression and TMB are poorly correlated (Yarchoan *et al*., 2019; Aurélien Marabelle *et al*., 2020; Sha *et al*., 2020). In addition, a retrospective analysis of MSI-h tumours found that TMB varies depending on whether MutS or MutL-related proteins had been lost, with MSH2/MSH6 deficient tumours having ∼46 mutations per megabase (mut/Mb) whilst MLH1/PMS2 deficient tumours had ∼25 mut/Mb (Salem *et al*., 2020). Therefore, this further contributes to the evidence that the MMRd/MSI-h cohort is a heterogeneous population.

Clinically, MMRd patients are grouped solely based on the loss of one or more of the four MMR genes and there is no further stratification based on which protein is lost. Therefore, this study aimed to investigate if loss of the individual MMR genes differentially impacted factors associated with ICI response, namely PD-L1 expression, TMB and MSI. This work highlights that loss of the different MMR genes may contribute to heterogenous response rates to ICIs in the MMRd/MSI-h cohort.

## MATERIALS & METHODS

### Cell lines

Cell lines were purchased from ATCC and grown at 37 °C and 5 % CO_2_ in a humidified atmosphere. U2OS, OVCAR4, and SW620 were grown in Dulbecco’s Modified Eagle Medium (DMEM; Gibco) whilst CT26 and SNU119 were grown in Roswell Park Memorial Institute 1640 (RPMI; Gibco) media. All media was supplemented with 10% fetal bovine serum (FBS; Invitrogen) and 100 U/ml penicillin and 100 μg/ml streptomycin (Gibco). The compounds cisplatin (Teva UK), IFNs (human IFNγ, Biolegend; murine IFNγ, Peprotech; IFNα, Abcam; IFNβ, Peprotech) and palbociclib (Selleck) were obtained commercially. To generate CRISPR-Cas9 guide control (gCTRL), MLH1 and MSH6 knock-out cell lines in the CT26 and U2OS, cells expressing Cas9-GFP were generated via lentiviral transduction and selected for with flow cytometry. Subsequently, the Dharmacon Edit-R system was used with predesigned CRISPR RNAs targeting MLH1 and MSH6. Single-cell clones were expanded, and screening was performed via western blot and targeted sequencing of the gRNA site. CT26 gCTRL, MLH1 KO and MSH6 KO cells were continuously passaged over 12 weeks, cryopreserving at weeks 1, 4, 8 and 12 in FBS + 10% DMSO. Cell populations from each time point could then be thawed simultaneously to allow for characterization and in MSH2 KO cells this has been shown to generate matched cell populations with MSI ranging from low to high. Unless otherwise stated, MMR KO cells were passaged for 12 weeks before characterisation to allow for the accumulation of mutations.

All cell lines were authenticated based on STR profile, viability, and morphologic inspection and were routinely mycoplasma tested.

### Protein analysis

Cell pellets were lysed in radioimmunoprecipitation assay (RIPA) buffer (50 nM Tris (pH 8), 150 nM NaCl, 1% NP40, 0.5% SDS) supplemented with protease inhibitors (Roche). For western blotting, lysates were electrophoresed on NuPAGE precast gels (Invitrogen) and immunoblotted with antibodies detailed in supplementary table 1. This was followed by incubation with anti-IgG-horseradish peroxidase and chemiluminescent detection (Supersignal West Pico Chemiluminescent Substrate, Pierce). Immunoblotting for β actin, β tubulin or vinculin (Cell signalling) was performed as a loading control. Protein densitometry was analysed using ImageJ and normalised to the loading control and control sample.

### Cell surface protein expression analysis

Cells were stained in suspension with an allophycocyanin-associated antibody targeting PD-L1 alongside an IgG2b, κ isotype control (Biolegend) on ice for 30 minutes. DAPI staining was also performed to exclude dead and apoptotic cells. Flow cytometry was performed using an LSR Fortessa 3-laser cell analyser (BD Biosciences). Unstained control samples were utilised to design the gating strategy applied to the experimental samples. Analysis was performed using FlowJo software (Tree Star) to quantify the relative cell surface expression by Median Fluorescence Intensity.

### qPCR

RNA was extracted from cells using the RNeasy kit (Qiagen). RNA was quantified using a nanodrop spectrophotometer (ThermoFisher) and 1 μg of RNA underwent reverse transcription using the High-Capacity cDNA Reverse Transcription kit (Applied Biosystems). Thermocycling was performed as follows: 25 °C for 10 mins, 37 °C for 120 mins then 85°C for 5 secs. 20 ng of RNA underwent qPCR using the TaqMan universal PCR master mix (Applied Biosystems) in the QuantStudio 5 system (Applied Biosystems). Samples were run in duplicate and thermocycling was carried out as follows: 50 °C for 2 mins, 95 °C for 10 mins then 40 cycles of 95 °C for 15 secs and 60 °C for 1 min. Analysis was performed using the 2 ^-ΔΔCT^ method normalised to β-actin or GAPDH.

### Cell cycle analysis

Cells were plated sparsely before treating with the CDK4/6 inhibitor palbociclib for 24 hours, which arrests the cells in G1. Cells were then collected and washed twice in PBS prior to fixing in 70% ethanol. Fixed cells were treated with RNase A to a final concentration of 50 μg/ml shaking at 37 °C for 30 minutes. The samples were then stained with propium iodide to a final concentration of 20 μg/ml for 30 minutes in the dark on ice before analysis with flow cytometry.

### Chromatin Immunoprecipitation

Cells were crosslinked for 10 minutes at room temperature (RT) using formaldehyde (1% in 50 mM HEPES, 100 mM NaCl, 1 mM EDTA, 0.5 mM EGTA) before quenching with glycine (1.25 M) for 5 minutes at RT. Crosslinked cells were collected before being successively resuspended and rotated in 2 different lysis buffers (50mM Hepes–KOH pH 7.5, 140 mM NaCl, 1 mM EDTA, 10% Glycerol, 0.5% NP-40, 0.25% Triton X-100 and 10 mM Tris–HCL, pH 8, 200 mM NaCl, 1 mM EDTA, 0.5 mM EGTA). Sonication was performed in 10mM Tris– HCl pH 8, 100 mM NaCl, 1 mM EDTA, 0.5 mM EGTA, 0.1% Na–Deoxycholate and 0.5% N-lauroylsarcosine for 20 minutes with a duty factor of 5% (Covaris S220). All buffers contained protease inhibitors (Roche). Chromatin shearing analysis was performed following treatment with RNase and Proteinase K (Thermo) in TE buffer via electrophoresis. Chromatin was cleaned up by centrifuging with Triton X-100 (10%) before being quantified using DS DNA broad range kit (QUBIT). Dynabeads Protein A were washed and blocked in ice-cold BSA-PBS (0.5%) before incubation with MSH6 antibody (ThermoFisher) alongside an IgG control overnight at 4 °C. Antibody-bound beads were washed in BSA-PBS (0.5%) before the addition of pre-cleared lysate in 1% Triton with protease inhibitor (Roche). Following rotation overnight at 4 °C the beads were washed in RIPA (50 mM HEPES pH 7.6, 1 mM EDTA, 0.7% Na deoxycholate, 1% NP-40, 0.5 M LiCl) and TE buffer. The beads were then resuspended in elution buffer (50 mM Tris-HCL pH 8, 10 mM EDTA, 1% SDS) overnight at 65 °C. RNase and proteinase K treatment was then performed before DNA was extracted using phenol-chloroform isoamyl alcohol and chloroform successively. Ethanol precipitation was used to purify DNA, and the pellet was resuspended in low EDTA TE buffer (10 mM Tris-HCl pH 8, 0.1 mM EDTA). 2 µl of immunoprecipitated DNA fragments in triplicate were quantified using SYBR green PCR master mix (Thermofisher) on the QuantStudio7 (Applied Biosystems). Primers designed by Wang *et al* 2020 targeting upstream of the PD-L1 gene start codon were used the sequences of which were as follows: −L1178Lbp to −L1117Lbp (forward 5′-GCT GGG CCC AAA CCC TAT T and reverse 5′-TTT GGC AGG AGC ATG GAG TT), −L455Lbp to −L356Lbp (forward 5′-ATG GGT CTG CTG CTG ACT TT and reverse 5′-GGC GTC CCC CTT TCT GAT AA-) and −L105Lbp to −L32Lbp (forward 5′-ACT GAA AGC TTC CGC CGA TT and reverse 5′-CCC AAG GCA GCA AAT CCA GT); hereafter referred to as primer 1, 2 and 3 respectively. Amplicons were between 60 and 150 base pairs. Results were normalised to the IgG using the 2 ^-ΔΔCT^ method.

### Whole Exome Sequencing

DNA was extracted from cells using the DNeasy blood and tissue kit (Qiagen) before whole exome sequencing (WES) was performed at Oxford Genomics (Wellcome) using Twist Library Preparation, yielding on average 115 million paired-end reads per sample with 150bp read length. After initial quality check using fastqc v0.11.5, reads were quality trimmed using trimgalore v0.6.5 and aligned to the mouse reference genome GRCm38 (mm10) using bwa v0.7.17. Picard v2.25.7 was used to mark read duplicates and assess insert size distributions. Base quality score recalibration was performed on known sites of variation using gatk v4.2.1.0 with mgp.v5.indels.pass.chr.sort.vcf.gz and 0-All.vcf.gz, downloaded from the Mouse Genome Project. Variant calling was performed using mutect2 (Benjamin et al. 2019). Treated samples were compared to their respective controls and variant call format files were filtered using gatk before further annotating with annovar. Further filtering was applied for mutations with sequencing depth DP > = 10. MSI analysis was performed using MSIsensor pro v1.2.0 (Niu et al. 2014). The number given is the percentage of microsatellite sites with a somatic indel and an MSIsensor score of 3.5 is considered MSI. Normal WES data from a BALB/c spleen was downloaded from the Sequence Read Archive (National Centre for Biotechnology Information; sample SRR7774027) and processed as described above. The bam file was used as “normal” for the MSIsensor analysis. Raw sequencing data have been deposited to the NCBI SRA database under PRJNA1186681.

### Immunofluorescence

Cells were seeded on poly-L-lysine pre-coated coverslips (Corning). Following treatment, cells were incubated for 1 minute in 0.1% Triton (Sigma) in 1X PBS. Coverslips were then fixed with 4% paraformaldehyde (PFA; Booster) with 2% sucrose (Sigma) in 1X PBS for 20 minutes. This was followed by three washes in 1X PBS. Incubation with γH2AX antibody (Millipore) was performed in 2% BSA at 37°C for 45 minutes and washed 3 times with 1X PBS. Coverslips were then incubated with mouse secondary antibody (Invitrogen) in 2% BSA at 37°C for 30 minutes or at RT for 1 hour in the dark and washed 3 times with 1X PBS. Coverslips were then stained with DAPI for 1 minute, washed twice with 1X PBS and mounted onto slides with ProLong Gold antifade reagent (Invitrogen). Images were captured with Zeiss 710 confocal microscope and processed with ImageJ.

### Cell viability

Cells were seeded in a 96 well plate prior to indicated treatment. Following this, Cell Titre Glow (Promega) reagent was diluted 1:4 in PBS before removal of the growth medium and addition to the cells. The plate was shaken for 2 minutes before being allowed to incubate for 10 minutes in the dark at RT. Cell viability was then quantified by luminescence on a FLUORstar Omega plate reader (BMG Labtech).

### Statistical analysis

Data represents standard error of the mean of at least three independent experiments. In most cases, 2-way ANOVA with a follow-up Tukey or Sidak test was used to determine statistical significance but where there were only 2 groups to compare unpaired student’s t-test was used as indicated in figure legends. The only exception to this is the MSIsensor data where a Fisher’s exact test was used with a follow-up Benjamini, Krieger and Yekutieli test to control for multiple testing (false discovery approach). Statistical analyses were performed using GraphPad Prism Software with p<0.05 regarded as significant.

## RESULTS

### MSH6 loss does not lead to increased PD-L1 expression

Given the clinical responses to ICIs are variable among MMRd patients and PD-L1 expression has previously been used as a marker of ICI response, we initially investigated whether loss of different MMR genes could result in differential PD-L1 expression. To this end, human ovarian cancer cells (OVCAR4) and human osteosarcoma cells (U2OS) were transfected with small interfering RNA (siRNA) targeting MLH1, PMS2, MSH2 and MSH6 to transiently deplete expression. As a positive control, the cells were treated with cisplatin to induce DNA damage as this induces PD-L1 expression (Sato *et al*., 2017). Induction of DNA damage was validated by performing γH2AX staining (Figure S1A,C). In addition, cell viability was assessed upon cisplatin treatment, to ensure that the majority of cells were still metabolically active despite DNA damage (Figure S1B,D). We observed that in cells depleted for MLH1, MSH2 or PMS2 there was increased expression of PD-L1 protein at both the basal level and following treatment with cisplatin (Figure 1A,C). However, PD-L1 expression was not induced in the MSH6-depleted cells. PD-L1 expression was also investigated at the cell surface level as this is where PD-L1 exerts its function. At the cell surface level, PD-L1 expression was significantly increased in the MLH1, MSH2 and PMS2 silenced cells in comparison to the MSH6 silenced cells following treatment with cisplatin in both OVCAR4 (Figure 1B; siMLH1, siMSH2 vs siMSH6 p<0.0005, siPMS2 vs siMSH6 p<0.01) and U2OS (Figure 1D; U2OS siPMS2, siMSH2 vs siMSH6 p<0.05) cell lines.

**Figure 1.**
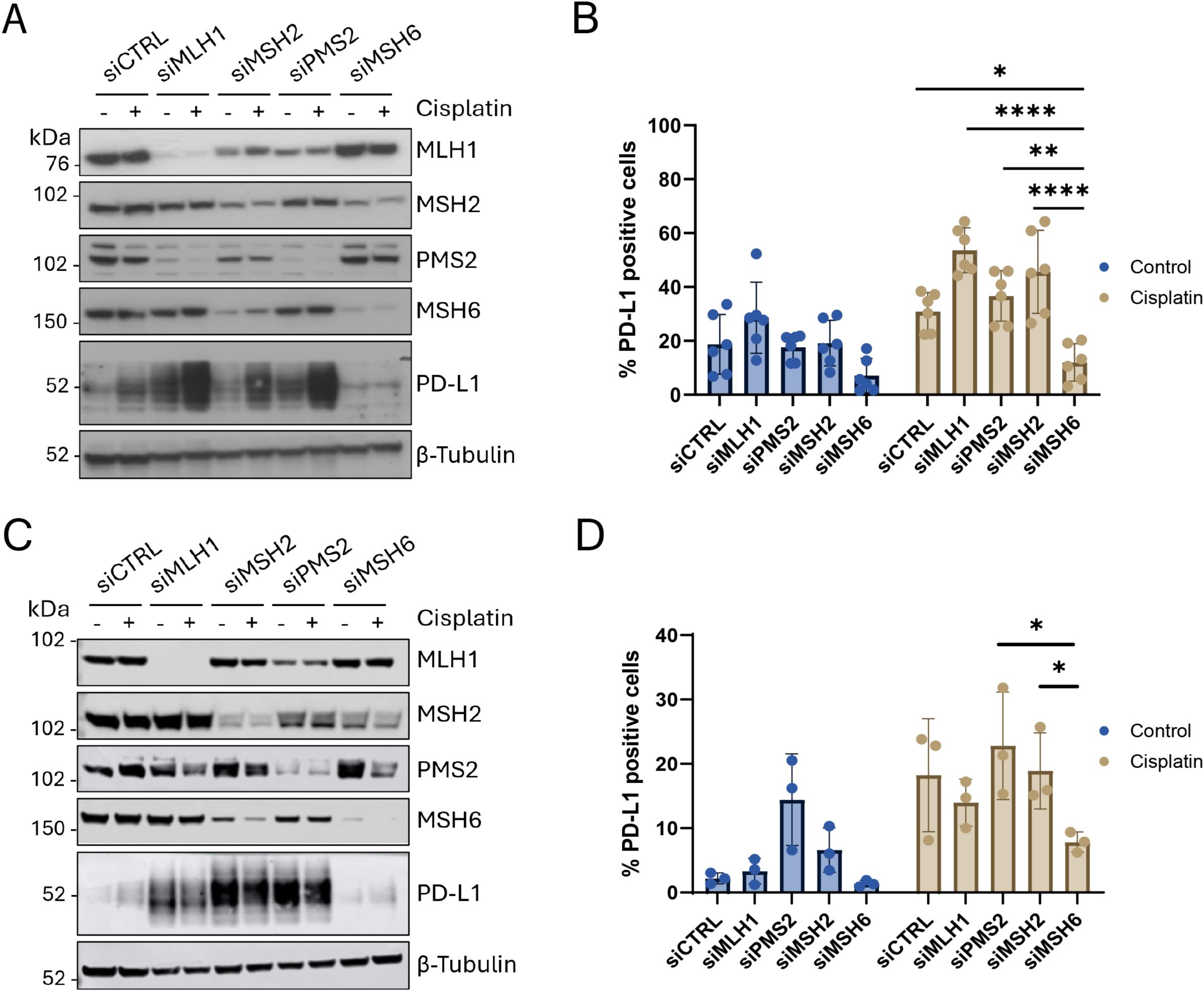
Depletion of MMR genes MLH1, MSH2 and PMS2, but not MSH6, results in increased expression of PD-L1. (A,B) OVCAR4 cells or (C,D) U2OS were transfected with siRNAs targeting each of the four MMR genes (MLH1, MSH2, PMS2 and MSH6) alongside a non-targeting siCTRL for 24 hours. The cells were then treated with cisplatin (2 μM) for 72 hours. (A,C) Western blot analysis to confirm knockdown of each MMR gene and measure PD-L1 expression. β tubulin was probed as a loading control. (B,D) Flow cytometry to measure PD-L1 expression. Analysis was performed using FlowJo with gating set up using an isotype control to determine the percentage of PD-L1 positive cells. Error bars represent (B) six or (D) three individual experiments. Statistical significance was carried out using two-way ANOVA with Tukey’s multiple comparison test (**** p<0.0001, ** p<0.01, * p<0.05).

To corroborate these findings and ensure this difference in PD-L1 expression was not an effect of siRNA transfection, CRISPR-Cas9 was utilised to knockout (KO) MLH1 and MSH6 in the murine CRC cell line CT26 (Figure S2) and in U2OS cells (Figure 2A). We observed that MLH1 KO increased basal PD-L1 expression relative to both the gCTRL and MSH6 KO cells (Figure 2A,B; p<0.01), therefore, consistent with siRNA knockdown, MSH6 KO failed to increase PD-L1 expression in U2OS cells. This was conserved in the murine CT26 cells where there was also higher PD-L1 expression in the MLH1 KO cells in comparison to the gCTRL and MSH6 KO cells (Figure S2).

**Figure 2.**
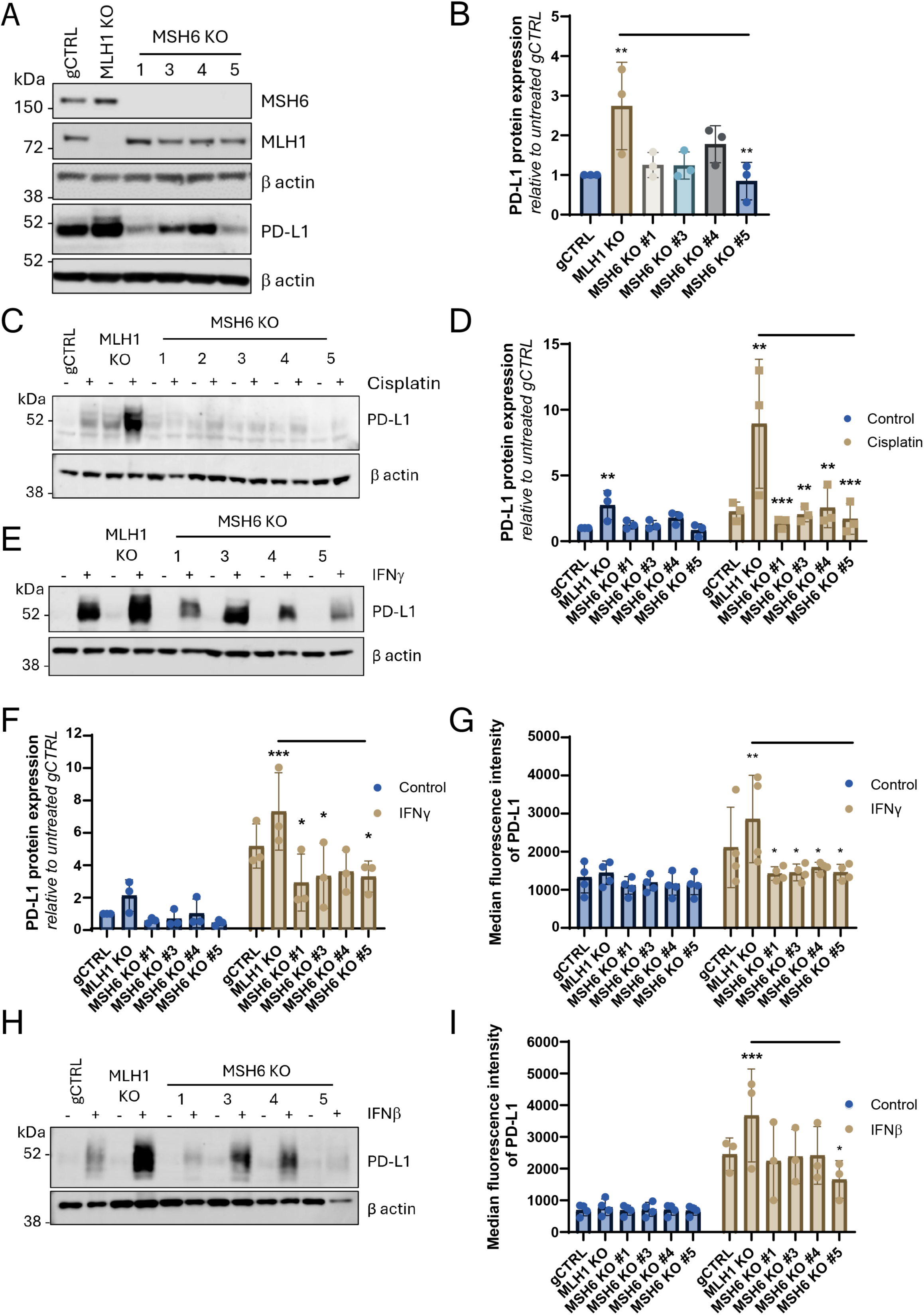
MSH6 is required for induction of PD-L1 expression following treatment with cisplatin and interferons. Cells were treated with (C,D) cisplatin (2 μM), (E-G) IFNγ (50 U/ml) or (H,I) IFNβ (100 u/ml) for 72 hours. Western blot analysis to (A) confirm knockout of MSH6 and MLH1 and (A,C,E) probe for PD-L1 expression. β actin was probed as a loading control. N=3. A representative blot is shown. (D,F) Protein densitometry was analysed using ImageJ. The intensity of PD-L1 expression was measured and normalised to β-actin. The normalised expression was then normalised to the untreated control cells. Error bars represent three individual experiments. Statistical significance was determined using 2-way ANOVA with Tukey’s multiple comparison test (***p<0.001, **p<0.01, no * = not significant). (G,I) Flow cytometry to measure PD-L1 expression. Analysis was performed using FlowJo with gating set up using an isotype control to determine the median fluorescent intensity of PD-L1 positive cells. Error bars represent (G) four or (I) three individual experiments. Statistical significance was determined using 2-way ANOVA with Tukey’s multiple comparison test and is in comparison to gCTRL unless otherwise shown (***p<0.001, **p<0.01, *p<0.05, no * = not significant).

### PD-L1 expression is not induced in MSH6-deficient cells upon DNA damage or IFN treatment

The induction of DNA damage is a known regulator of PD-L1 expression (Sato *et al*., 2017). Therefore, we next treated MLH1 and MSH6 KO cells with the alkylating agent cisplatin, to induce DNA damage and, observed that whilst there was a significant induction of PD-L1 protein expression in the MLH1 KO U2OS cells (Figure 2C,D; p<0.01), this induction was not observed in the MSH6 KO cells. In addition, in the cisplatin-treated samples, PD-L1 expression was significantly reduced in the MSH6 KO cells when compared to the MLH1 KO cells (Figure 2C,D; MSH6 KO #1 and #5 p<0.005, MSH6 KO #3 and #4 p<0.01). Similarly, CT26 MSH6 KO cells failed to induce PD-L1 expression following treatment with cisplatin (Figure S3A). Therefore, consistent with what was observed upon siRNA transfection there was a lack of induction upon PD-L1 expression in MSH6-deficient cells, which was conserved between mouse and human.

Next, we wanted to investigate whether these differences in PD-L1 expression were specific to DNA damage. To this end, treatment with IFNs was utilised as an alternative method to induce PD-L1 expression (Garcia-Diaz *et al*., 2017). We observed that IFNγ significantly induced PD-L1 expression in the MLH1 KO cells at both the protein and cell surface level (Figure 2E-G; protein p<0.005, cell surface p<0.01). However, upon treatment with IFNγ in MSH6 KO cells, there was a failure to induce PD-L1 expression to the same extent resulting in significantly less PD-L1 protein expression in comparison to the MLH1 KO cells (Figure 2F,G; protein MSH6 KO #1,3,5 p<0.05, cell surface p<0.05).

In cells silenced for MLH1, increased PD-L1 induction was observed following both IFNα and IFNβ treatment in comparison to the MSH6 KO cells (Figure S3C,D). This was further validated in the MLH1 KO cells where we observed that PD-L1 protein and cell surface expression was more strongly induced than in the gCTRL and MSH6 KO cells (Figure 2H,I; p<0.005). Furthermore, the IFNβ-treated MSH6 KO cells had significantly lower PD-L1 expression at the cell surface in comparison to the MLH1 KO cells (MSH6 KO #5 p<0.05).

Taken together, our data suggests that failure to induce PD-L1 expression to the same extent upon MSH6 loss in comparison to upon loss of other MMR proteins is not related to DNA damage but may define a non-canonical role for MSH6 in the regulation of PD-L1 expression.

### Differential PD-L1 expression does not correlate with TMB or JAK/STAT signalling

To determine whether PD-L1 expression correlated with MSI and TMB, whole exome sequencing was carried out on CT26 MSH6 KO and MLH1 KO cells, which were serially passaged over 12 weeks in parallel with gCTRL cells to allow the accumulation of mutations (Mandal *et al*., 2019). In MLH1 KO matched cell populations from weeks 1, 4 and 12, we observed that MSI increased over time so that at week 12 there was significantly more MSI in comparison to gCTRL cells (Figure 3A, p<0.0001). This is consistent with what has been previously observed in MSH2 KO cells (Mandal *et al*., 2019). However, in the MSH6 KO cells, there was no increase in MSI over time, such that at week 12, MSI was significantly higher in the MLH1 KO in comparison to the MSH6 KO cells (Figure 3A, p<0.0001). Interestingly, this suggests that MSI may correlate with PD-L1 expression. To investigate whether MSI correlated with TMB, the total number of mutations compared to the gCTRL was quantified. Surprisingly, we observed that MSH6 KO cells had significantly higher TMB than MLH1 KO cells (Figure 3B, p<0.05), and this did not increase over time. To understand this further we analysed the different types of mutations generated and observed that no significant differences were observed between all indels or frameshift indels when comparing the MLH1 KO and MSH6 KO cells (Figure 3C,D). However, MSH6 KO cells had significantly more nonsynonymous single nucleotide variants (SNVs) than the MLH1 KO cells (p<0.05, Figure 3E).

**Figure 3.**
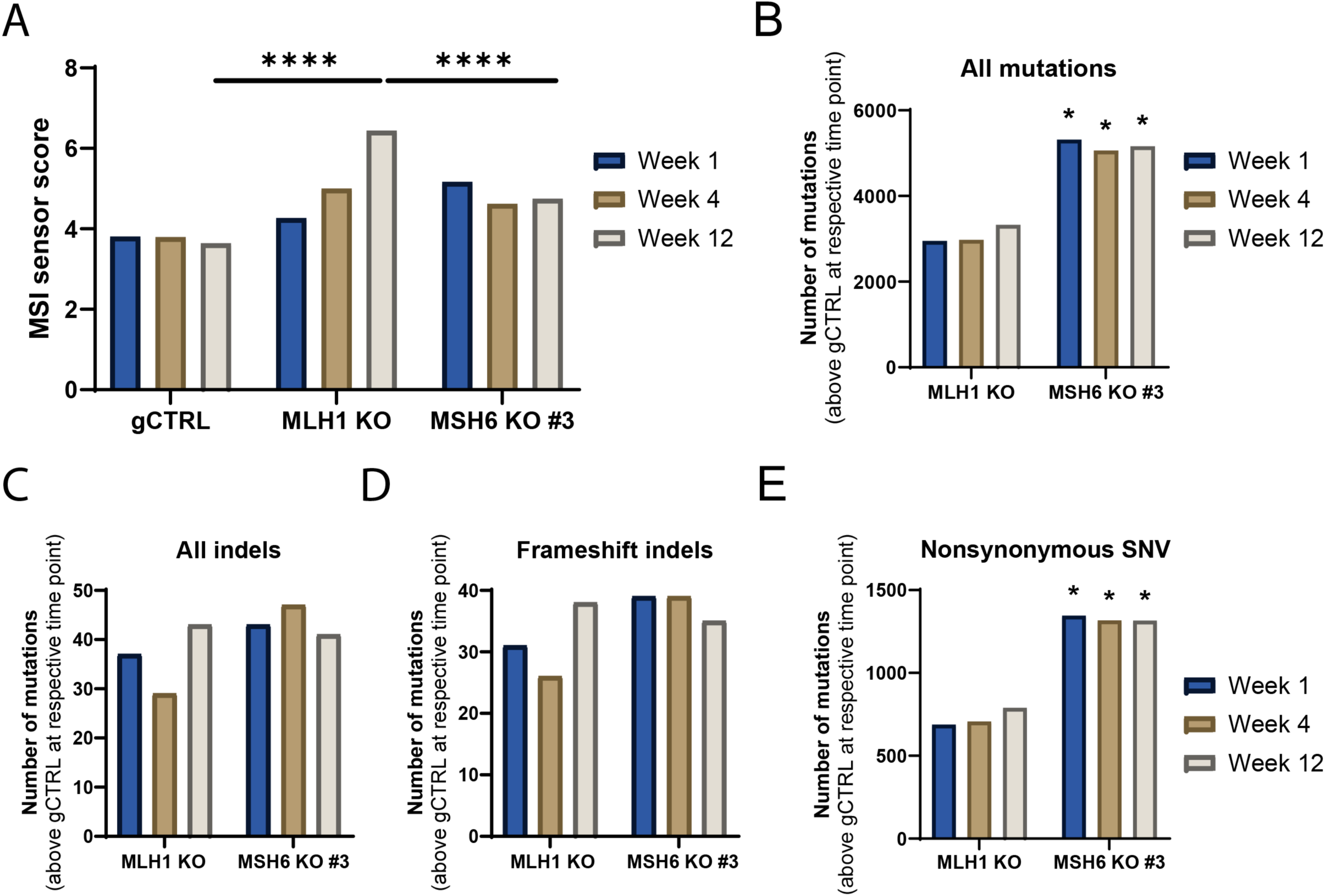
PD-L1 expression, MSI and TMB do not correlate in MSH6 KO cells. DNA was extracted and sent to Oxford Genomics for whole exome sequencing before analysis was carried out to determine (A) MSI using MSIsensor and (B-E) TMB using Mutect2. (A) Statistical analysis was determined using Fisher’s exact test with a follow-up Benjamini, Krieger and Yekutieli test to correct for multiple testing (****p<0.0005). (B-E) Statistical analysis was determined using 2-way ANOVA with Tukey’s multiple comparison test (MSH6 KO vs. MLH1 KO; *p<0.05, no * = not significant).

JAK/STAT signalling is central to PD-L1 regulation in response to DNA damage and IFNs (Garcia-Diaz *et al*., 2017; Sato *et al*., 2017). Therefore, to determine whether these differences in PD-L1 expression upon MLH1 and MSH6 loss were mediated by JAK/STAT signalling the expression and phosphorylation of STAT1 and STAT3 were investigated. The U2OS MLH1 KO and MSH6 KO cells were treated with cisplatin (Figure 4A) and IFNγ (Figure 4B) to activate STAT1/3 phosphorylation. We observed that there were no differences in the expression or phosphorylation of STAT1/3 upon MSH6 loss. Despite this, decreased expression of STAT1 as well as expression and phosphorylation of STAT3 was observed in the CT26 MSH6 KO cells in comparison to the MLH1 KO cells (Figure S4). However, as this was not observed in the U2OS cells we concluded that this was not the conserved mechanism mediating the reduced PD-L1 expression observed in MSH6 KO cells. As these differences were not observed to be specific to DNA damage and instead indicated an inability to induce PD-L1 expression in MSH6 deficient cells, the possibility that MSH6 could directly regulate PD-L1 expression was investigated.

**Figure 4.**
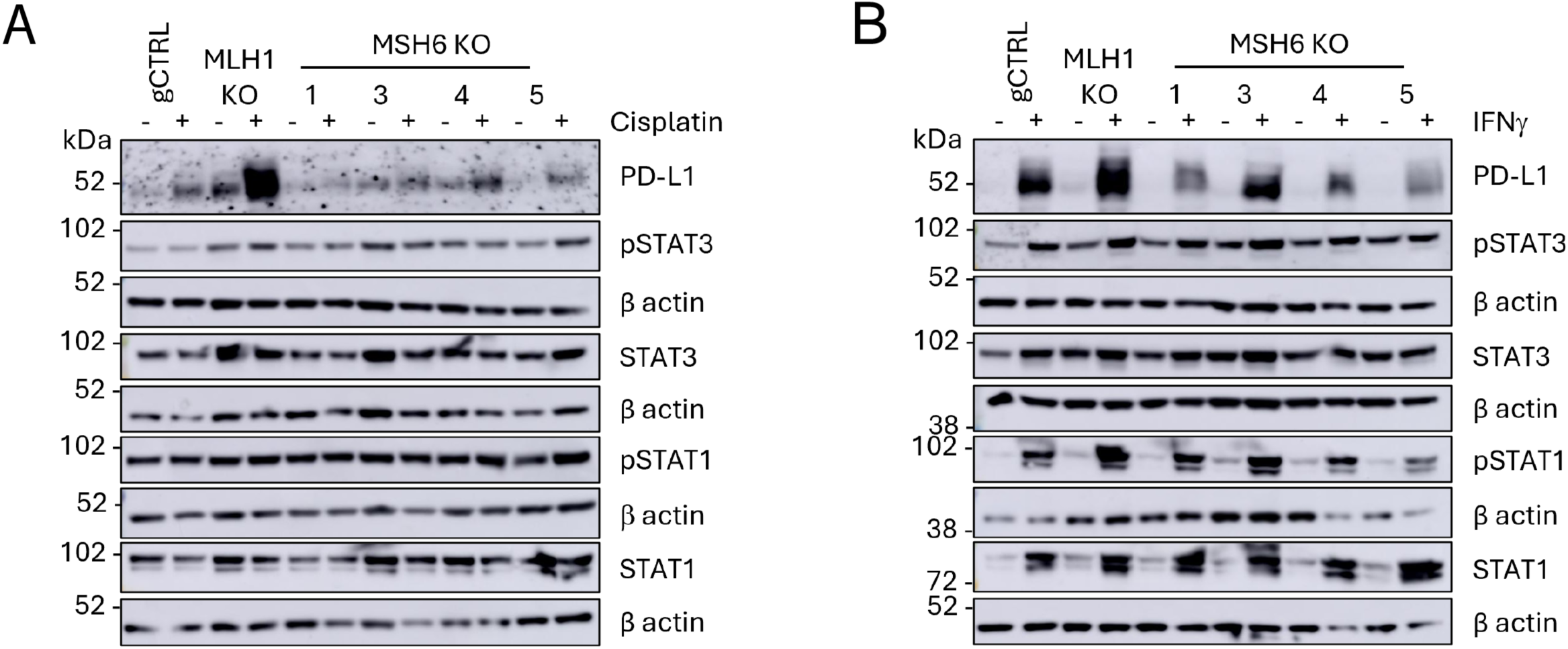
JAK-STAT signalling is not differentially regulated upon loss of MLH1 and MSH6. U2OS gCTRL, MLH1 KO and MSH6 KO were treated with (A) cisplatin (2 μM) or (B) IFNγ (50 U/ml) for 72 hours. Western blot analysis was performed to probe for PD-L1 as well as STAT1/3 and their phosphorylated form (S727), which is required for the full transcriptional activity and biological function of STAT1/3. β actin was probed as a loading control. N=3. A representative blot is shown.

### MSH6 can bind directly to the PD-L1 promoter

To determine whether MSH6 was binding directly to the PD-L1 promoter in the U2OS gCTRL, MLH1 KO and MSH6 KO cells, we performed chromatin immunoprecipitation (ChIP) by immunoprecipitating MSH6 coupled with qPCR utilising primers spanning the PD-L1 promoter. Our analysis revealed that there was significant binding of MSH6 to the PD-L1 promoter in gCTRL cells (Figure 5A; primer 2 p<0.05). Interestingly, in MLH1 KO cells statistically significant binding of MSH6 to the PD-L1 promoter was also observed (Figure 5B; p<0.05). This was also performed in MSH6 KO cells to act as a negative control and no binding was observed (Figure 5C).

**Figure 5.**
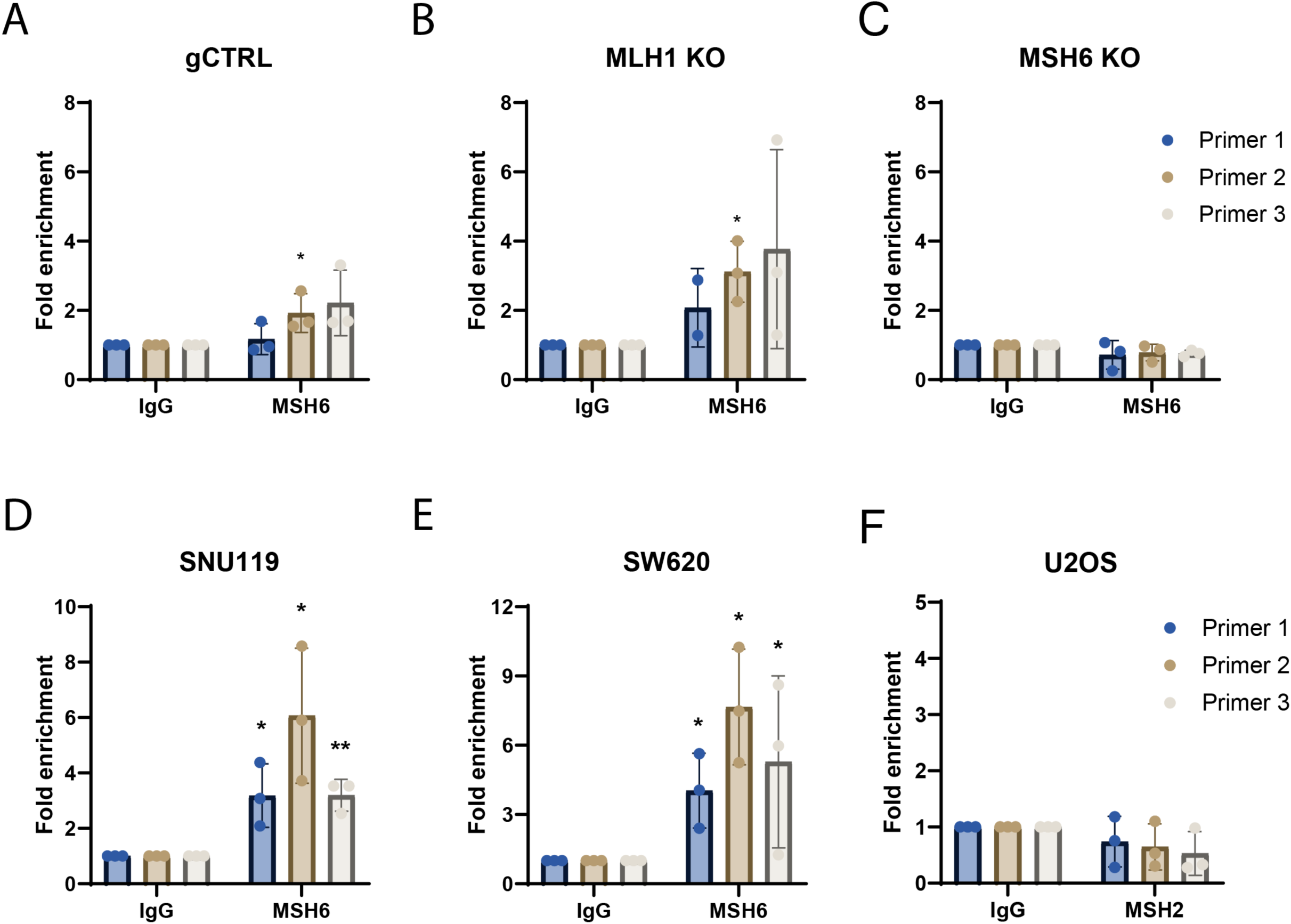
MSH6 binds directly to the PD-L1 promoter. Chromatin immunoprecipitation was performed using anti-IgG and anti-MSH6 conjugated beads on crosslinked and sonicated chromatin from U2OS (A) gCTRL, (B) MLH1 KO (C) MSH6 KO and (F) wildtype cells as well as (D) SW620 and (E) SNU119 cells. Primers targeting the PD-L1 promoter were then used in qPCR to quantify pulled-down DNA. Results are represented as fold change compared to the IgG. Error bars represent three individual experiments. Statistical analysis was determined using an unpaired t-test (**p<0.01, *p<0.05, no * = not significant).

The binding of MSH6 to the PD-L1 promoter was further validated in two additional human cancer cell lines, including SW620 (colorectal cancer) and SNU119 (ovarian cancer). In SW620 cells, we observed significant binding of MSH6 to the PD-L1 promoter in all three primer regions (Figure 5D, p<0.05). This was also true in the SNU119 cells (Figure 5E; primer 1,2 p<0.05; primer 3 p<0.01). Therefore, our results strongly suggest that the binding of MSH6 to the PD-L1 promoter is a conserved phenotype.

To determine whether MSH6 binds to PD-L1 as part of the MutSα heterodimer, we carried out ChIP analysis by immunoprecipitating MSH2 followed by qPCR utilising primers spanning the PD-L1 promoter. However, we did not observe MSH2 binding to the PD-L1 promoter (Figure 5F). This suggests that MSH6 is not bound as part of the MutSα (MSH2-MSH6) complex.

### MSH6 can bind to the PD-L1 promoter in a SETD2-dependent manner

To further understand the role of MSH6 in the regulation of PD-L1 expression, we next investigated which phase of the cell cycle MSH6 was bound to the PD-L1 promoter. The canonical role of MSH6 in MMR occurs during S phase to repair DNA replication errors (Kunkel and Erie, 2005). Therefore, the binding of MSH6 to the PD-L1 promoter in G1 would suggest a role in transcription instead of MMR. To this end, U2OS cells were arrested in G1 phase using the cyclin-dependent kinase (CDK)4/6 inhibitor, palbociclib (Figure 6A) followed by ChIP analysis of MSH6 at the PD-L1 promoter. We observed significant binding of MSH6 in both the WT asynchronous cells (Figure 6B; primer 1 p<0.05) and the G1-arrested cells (Figure 6C; primer 2 p<0.01).

**Figure 6.**
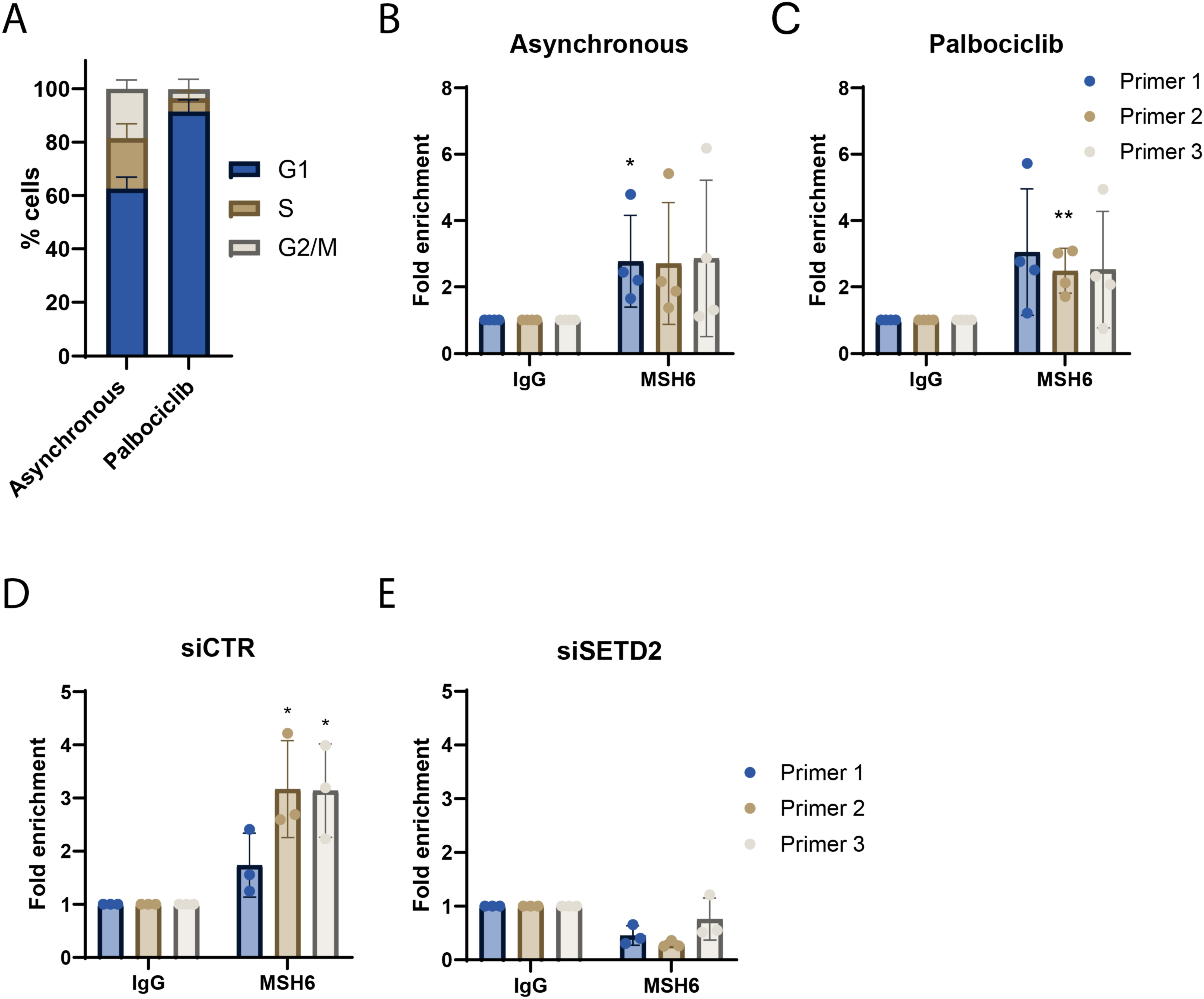
MSH6 binds to the PD-L1 promoter in a SETD2-dependent manner in G1. (A-C) Cells were arrested in G1 using the CDK4/6 kinase inhibitor palbociclib and (A) cell cycle analysis was performed using propidium iodide staining, which was quantified using flow cytometry to confirm arrest. (D,E) U2OS cells were transfected with siRNA targeting SETD2 alongside a non-targeting control (siCTRL). (B-E) Chromatin immunoprecipitation was performed using anti-IgG and anti-MSH6 conjugated beads on crosslinked and sonicated chromatin from U2OS wild-type cells. Primers targeting the PD-L1 promoter were then used in qPCR to quantify pulled-down DNA. Results are represented as fold change compared to the IgG. Error bars represent (A,D,E) three or (B,C) four individual experiments. Statistical analysis was determined using an unpaired t-test (**p<0.01, *p<0.05, no * = not significant).

Thus far, we have demonstrated that MSH6 is required for induction of PD-L1 expression and that it binds to the PD-L1 promoter during G1, suggesting a role in transcription. However, given no previous role for MSH6 in transcription has been defined, the mechanism of recruitment is unclear. MutSα can be recruited to chromatin via the trimethyl histone mark on lysine 36 of histone 3 (H3K36me3), which is made by the histone trimethylase SETD2 (X. Li *et al*., 2013). This histone mark can be bound by the Proline-Tryptophan-Tryptophan-Proline (PWWP) domain in MSH6. We hypothesised that this could also be the mechanism of recruitment for MSH6 to the PD-L1 promoter. To investigate this, we silenced SETD2 in U2OS cells before performing ChIP analysis to determine if MSH6 binding to the PD-L1 promoter was dependent on SETD2. We observed that upon siRNA-mediated silencing of SETD2, there was no binding of MSH6 to the PD-L1 promoter (Figure 6E) whilst in the siCTRL transfected cells significant binding was observed (Figure 6D; primer 2,3 p<0.01). Therefore, this suggests that the histone mark H3K36me3, which is made by SETD2 and can be bound by the PWWP domain at the N terminus of MSH6, is involved in the recruitment of MSH6 to the PD-L1 promoter.

Overall, our data suggests that MSH6 deficient cells fail to induce PD-L1 expression at the basal level and to the same extent as upon loss of other MMR proteins in response to DNA damage and IFN treatment. We have shown that this is due to the ability of MSH6 to bind directly to the PD-L1 promoter in a SETD2-dependent manner to regulate PD-L1 transcription.

## DISCUSSION

Since the approval of MMRd/MSI-h as a biomarker for response to pembrolizumab, the potential for ICIs to effectively treat MMRd/MSI-h tumours has been demonstrated in many studies (André *et al*., 2020; Aurelien Marabelle *et al*., 2020; Le *et al*., 2020; Cercek *et al*., 2022). However, ∼40-70% of MMRd/MSI-h patients remain refractory to ICI treatment, highlighting the inadequacy of this biomarker alone in predicting patients likely to respond to ICIs (Mestrallet *et al*., 2023). This variability in response is not only true for the MMRd/MSI-h cohort but also for the other approved biomarkers for predicting response to ICIs, namely TMB and PD-L1 expression (McGrail *et al*., 2021; Wu *et al*., 2022; Sun *et al*., 2023). It’s also important to consider that these three approved biomarkers for predicting response to ICIs, although biologically interlinked, do not necessarily correlate in patients (Vanderwalde *et al*., 2018). There is context dependency to this, for example, PD-L1 expression and high TMB are significantly associated in gastric and endometrial cancers, which may explain why ICIs are much more successful in some cancer types than others (Yarchoan *et al*., 2019). Therefore, it is evident that more work is required to understand the biological mechanisms promoting response to ICIs to allow for better patient stratification in the clinic. Hence in this study, MSI, TMB and PD-L1 expression were evaluated in MMRd cells to identify heterogeneity and elucidate the mechanism behind this.

Pembrolizumab, the approved ICI to treat MMRd/MSI-h patients, targets the PD-1/PD-L1 interaction, and PD-L1 expression on tumour cells has been approved as a biomarker for predicting response to ICIs in some cancer types, such as NSCLC (Pai-Scherf et al. 2017). This was based on the KEYNOTE-024 phase III trial, which found that in advanced NSCLC, pembrolizumab as a first-line treatment in patients with >50% of PD-L1 positive tumour cells achieved an improved overall survival (5.2 months vs 4.1 months) and ORR (44.8% vs. 27.8%) in comparison to platinum-based chemotherapy (Reck *et al*., 2016). However, another phase III trial observed prolonged survival in patients treated with anti-PD-1 instead of chemotherapy regardless of PD-L1 expression (Brahmer *et al*., 2015). Therefore, although tumoral PD-L1 expression is not intrinsically linked with response to ICIs in that it is not sufficient for response, it has been associated with response to ICIs. We have demonstrated that MSH6 deficient cells fail to induce PD-L1 expression and that this is due to the ability of MSH6 to directly regulate PD-L1 expression. We also observed that MSH2 was not bound to the PD-L1 promoter, which suggests that MSH6 is not acting as part of MutSα. This may explain why we see reduced PD-L1 expression in MSH6 deficient cells but not in MSH2 deficient cells, even though MSH2 and MSH6 heterodimerise to perform their function in MMR. The differences in PD-L1 induction in this context, in combination with the evidence that PD-L1 expression can contribute to response to ICIs, may suggest that gene-specific stratification of MMRd patients would be beneficial for predicting response to ICIs. However, it is important to note that PD-L1 can also be expressed on immune cells, and it has been shown that quantifying PD-L1 on all cells in the tumour microenvironment, rather than just on the tumour cells, can better predict response to pembrolizumab (Zhu *et al*., 2018). Therefore, it would be beneficial to evaluate PD-L1 expression on all cells in the TME upon loss of each of the MMR genes.

We have also shown that MSH6 recruitment to the PD-L1 promoter is dependent on SETD2, a histone trimethyltransferase that has roles in alternative splicing, chromatin architecture organisation and homologous recombination (Aymard *et al*., 2014; Fahey and Davis, 2017). Interestingly, SETD2 deficient cells have many characteristics of MMRd cells, including high TMB and MSI, and SETD2 dysfunction has been associated with improved response to ICIs independent of TMB (Li et al., 2013; Weinstock et al., 2017; Battaglin et al., 2023; Zheng et al., 2023). This phenotype is likely caused at least in part by an impairment of MMR due to an inability to recruit MutSα to the chromatin via the PWWP domain on MSH6 (F. Li *et al*., 2013). Here, we have shown that MSH6, but not MSH2, is observed binding to the PD-L1 promoter in a SETD2-dependent manner in G1, which points more towards a role in transcription. Therefore, this suggests that SETD2 may also have a role in regulating transcription that may warrant further investigation.

TMB has also received separate site agnostic approval for treatment with pembrolizumab following the KEYNOTE-158 trial, where high TMB tumours (>10 mut/Mb) achieved an ORR of 29% compared to 6% in the low TMB group (<10 mut/Mb) (Marabelle et al 2020). Yet with studies showing that ∼60-85% of high TMB patients do not achieve an ORR following treatment with ICIs (McGrail *et al*., 2021), it is evident that although mutational load is associated with response to ICIs, it is not sufficient to elicit response. It has been suggested that instead of overall TMB being key, it’s more a matter of ‘quality over quantity’ with certain mutations, such as indels, evoking a stronger immune response than other types of DNA damage (Turajlic *et al*., 2017). This is consistent with the data that has suggested that response to ICIs in MMRd patients correlates with the degree of MSI (Mandal *et al*., 2019). This is also in line with the better response rates observed in MMRd/MSI-h patients when compared to the more general TMB-h, as MSI results in indels and, therefore, more immunogenic mutations. Interestingly, even though we observed that MSH6 KO cells had a higher TMB in comparison to MLH1 KO cells, this was primarily comprised of SNVs. Excitingly, this is consistent with a recent study that demonstrated that MSH6-deficient CRCs have significantly higher MMRd signature-associated SNVs than PMS2-deficient CRCs (Helderman et al., 2023). Therefore, this suggests that our models are representative of what is observed in the clinic. The accumulation of SNVs in MSH6 deficient cells could be because MutSα is responsible for recognising SNVs whilst MutSβ is responsible for recognising larger indels (Genschel *et al*., 1998). Therefore, if it is ‘quality over quantity’ when it comes to mutations driving response to ICIs and the mutational profiles of MMRd/MSI-h patients are not homogenous, this could be driving variable response rates to ICIs in this patient cohort. Similarly, some other studies have noted that MSH6 deficient tumours have lower MSI or are actually microsatellite stable (Wu *et al*., 1999; Berends *et al*., 2002; Hendriks *et al*., 2004; Plaschke *et al*., 2004; Latham *et al*., 2019). We observed that in MSH6 KO cells, MSI did not increase over time and that in week 12 cells MLH1 KO cells had significantly higher MSI than MSH6 KO cells. Our data also suggests that MSI but not TMB correlates with PD-L1 expression as MSH6 KO cells have a higher TMB but not MSI when compared to MLH1 KO cells. Therefore, if the degree of MSI is heterogeneous within the MMRd population then the classification of MMRd and MSI-h as a single patient cohort could be detrimental to identifying patients that are going to derive clinical benefit from ICIs.

Interestingly, beyond response to ICIs there is accumulating evidence that demonstrates that loss of the different MMR genes can result in distinct phenotypes. This includes the fact that noncanonical roles for MMR proteins have been identified, for example, MLH1 has been implicated in regulating mitochondrial metabolism with loss of MLH1 resulting in reduced basal oxygen consumption rate and reduced spare respiratory capacity (Rashid *et al*., 2019). In addition, particularly relevant to this work, a recent study identified stochastic switching between activation and inactivation of MSH6 and MSH3 as a driver of intratumoural heterogeneity, with loss of MSH6/MSH3 resulting in a hypermutator phenotype that drives subclonal evolution at the cost of cell fitness (Kayhanian *et al*., 2024). This allows the acquisition of traits that allow immune evasion and clone expansion, followed by a reversion in the inactivation of MSH6/MSH3 that stabilises the mutation rate to restore cell fitness. Other non-canonical roles for MutSα and MutSβ have also been identified in 5-methylcytosine deamination or oxidative damage via interaction with partners other than MutL, such as MBD4 (Fang *et al*., 2021; Zou *et al*., 2021). Alongside this, the evidence presented in this study suggests that MSH6 can also function as a transcription factor, so further work may be warranted to determine whether MSH6 can influence the transcription of other proteins. Further evidence that loss of different MMR genes can result in distinct phenotypes can be derived from Lynch syndrome, which is caused by germline mutations in any of the MMR genes. Lynch syndrome is an autosomal condition that increases a patient’s risk of colorectal and endometrial cancer to ∼80% and 40%, respectively (Aarnio *et al*., 1995). Interestingly, it is a highly heterogeneous patient population with varying penetrance of cancer, age of onset and molecular profiles, with recent data from the Prospective Lynch Syndrome Database proposing gene-specific stratification of patients (Dominguez-Valentin *et al*., 2020; Helderman *et al*., 2021; Møller *et al*., 2023). Therefore, perhaps this should also be extended to predicting responses to ICIs.

In summary, the variability in response to ICIs that is observed in the MMRd/MSI-h population may be derived from the loss of the different MMR genes. Here, we have shown that MSH6 KO cells have a lower expression of PD-L1 and an inability to induce expression to the same extent as other MMRd cells. We propose that this is due to the ability of MSH6 to bind directly to the PD-L1 promoter in a SETD2-dependent manner, defining a novel role for MSH6 as a transcription factor. In addition, we have demonstrated that whilst MSI is lower in MSH6 deficient cells in comparison to MLH1 deficient cells, the reverse is true for TMB. Therefore, this study contributes to the accumulating evidence that gene-specific stratification of MMRd patients could be therapeutically and clinically beneficial.

## ACKNOWLEDGEMENTS

We would like to thank the CRUK flow cytometry, microscopy and bioinformatics core facilities as well as all members of the Martin and Sharp labs for helpful discussions. This work was supported by funding from the Barts Charity (G-002509), Cancer Research UK (CRUK; C355/A25137) and CRUK Major Centre Award (A18066),

