## Supplementary data for "The DNA Mismatch repair protein, MSH6 is a novel regulator of PD-L1 expression"

**Supplementary table 1. List of antibodies used.**

| **Target** | **Mono/polyclonal** | **Reference** | **Antibody raised in** | **Source** | **Dilution** |
| --- | --- | --- | --- | --- | --- |
| PD-L1 | Monoclonal | E1L3N | Rabbit | Cell Signalling Technology | 1:500 |
| STAT1 | Monoclonal | D1K9Y | Rabbit | Cell Signalling Technology | 1:1000 |
| STAT1 pS727 | Monoclonal | D3B7 | Rabbit | Cell Signalling Technology | 1:1000 |
| STAT3 | Monoclonal | D3Z2G | Rabbit | Cell Signalling Technology | 1:1000 |
| STAT3 pS727 | Monoclonal | 9134 | Rabbit | Cell Signalling Technology | 1:1000 |
| MLH1 | Monoclonal | D38G9 | Rabbit | Cell Signalling Technology | 1:1000 |
| PMS2 | Polyclonal | C-20 | Rabbit | Santa Cruz | 1:1000 |
| MSH2 | Monoclonal | D24B5 | Rabbit | Cell Signalling Technology | 1:1000 |
| MSH6 | Polyclonal | P150 | Rabbit | Cell Signalling Technology | 1:1000 |

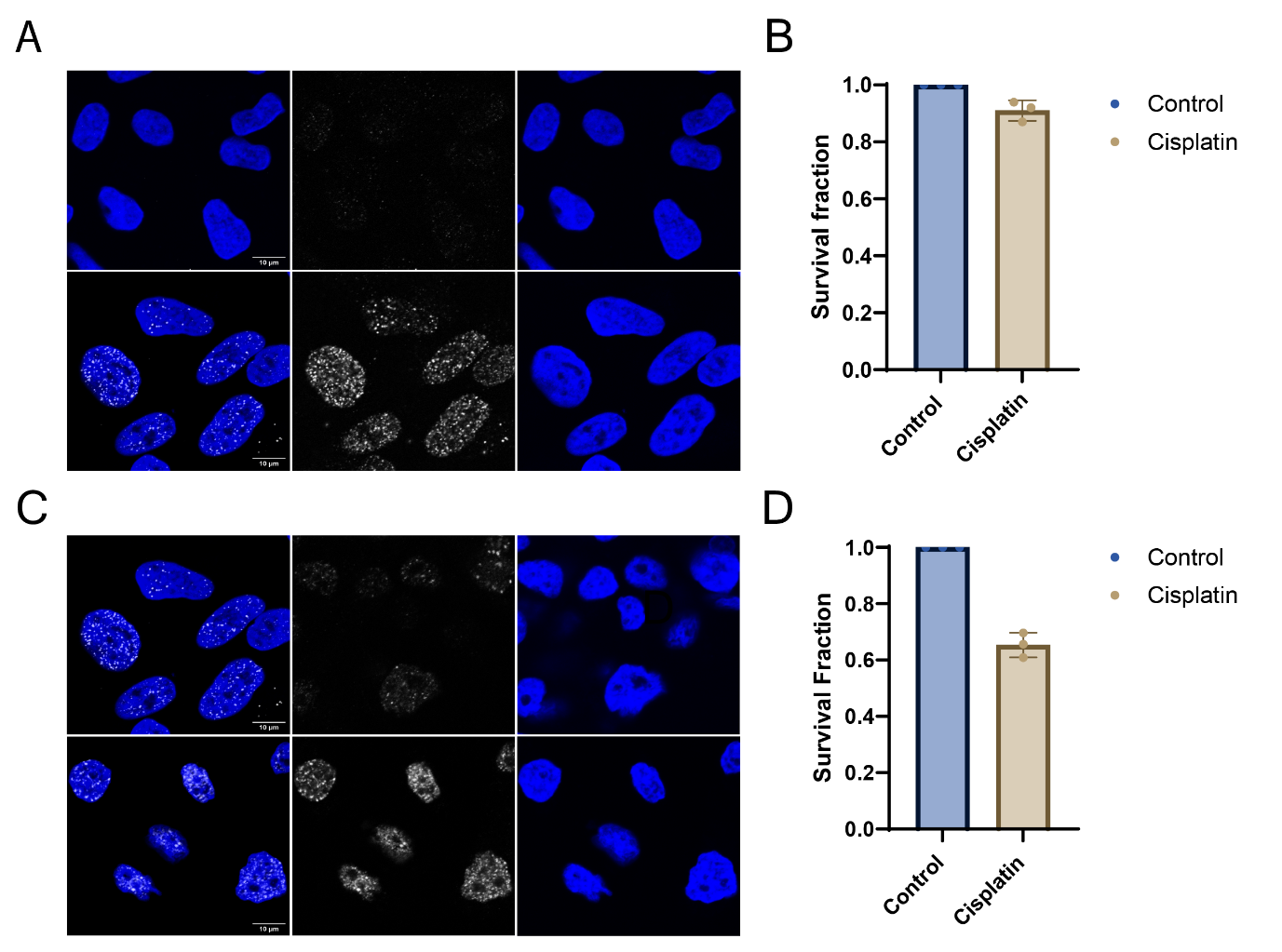

**Supplementary Figure 1. Cisplatin induces gH2AX formation in U2OS and OVCAR4 cells, but the majority of cells remain viable.** (A) U2OS and (C) OVCAR4 cells were seeded on poly-L-lysine coated coverslips prior to treatment with 2 μM cisplatin for 72 hours. Cells were fixed with 4% paraformaldehyde, stained with anti-gH2AX antibody and DAPI, and mounted on microscope slides. Images were acquired on LSM710 confocal microscope. N=3. Representative images are shown. (B) U2OS and (D) OVCAR4 cells were treated with 2 μM cisplatin for 72 hours prior to a Cell Titre Glow assay being performed. N=3.

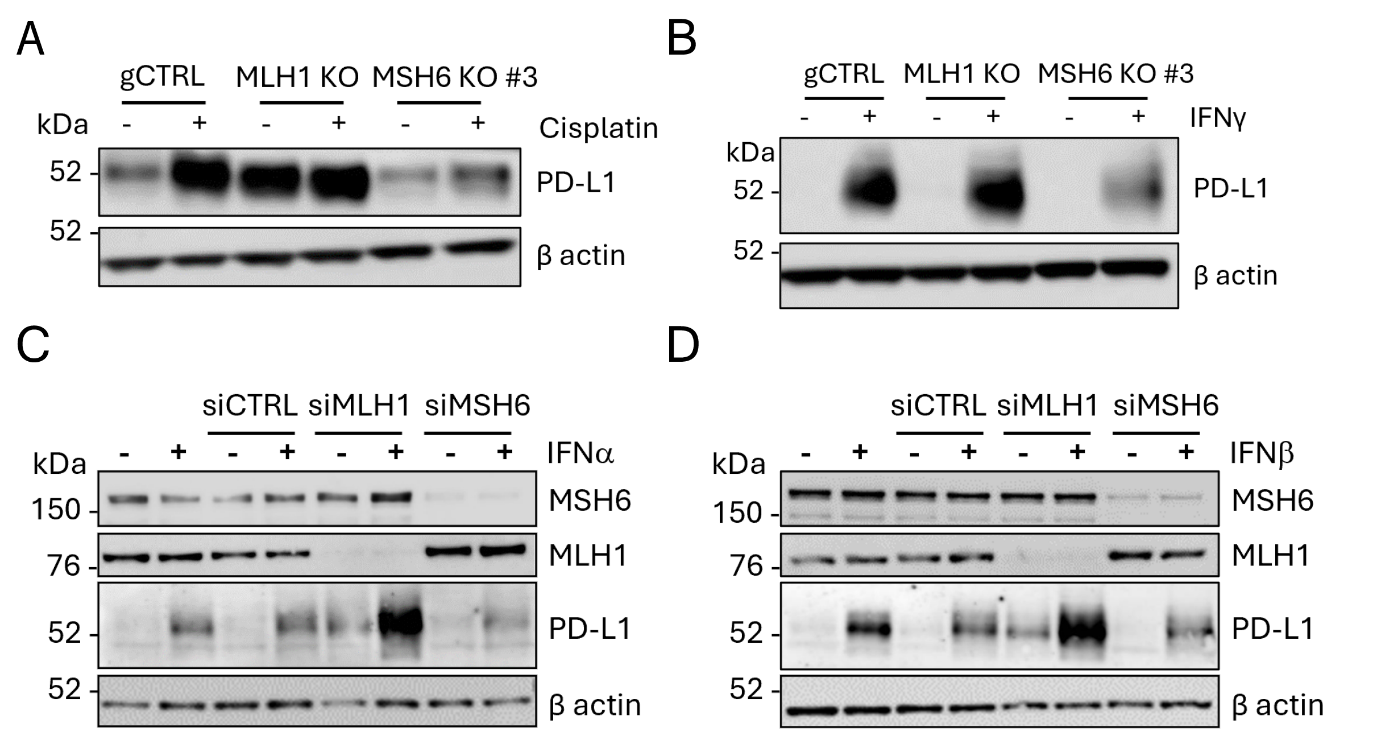

**Supplementary Figure 2. CT26 MLH1 KO cells have higher PD-L1 expression than MSH6 KO cells.** Western blot analysis was performed on CT26 gCTRL, MLH1 KO and MSH6 KO cells to probe for MLH1, MSH6 and PD-L1. β actin was probed as a loading control. N=3. A representative blot is shown.

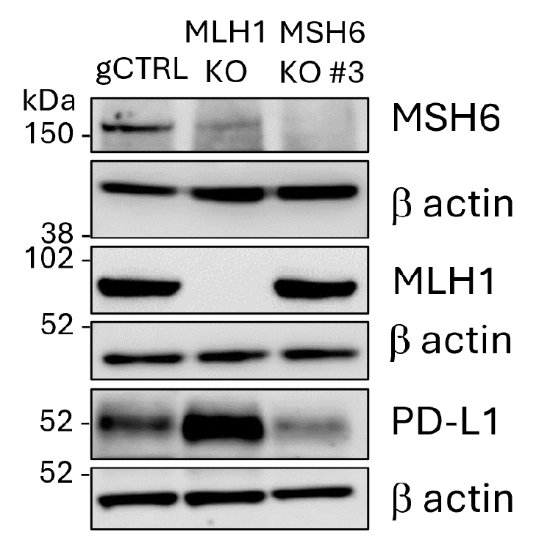

**Supplementary figure 3. MSH6 deficient cells fail to induce PD-L1 expression to the same extent as MLH1 deficient cells.** (A,B) CT26 gCTRL, MLH1 KO and MSH6 KO cells were treated with (A) cisplatin (2 μM) or (B) IFNγ (50 U/ml) for 48 hours before performing western blot analysis to blot for PD-L1. β actin was probed as a loading control. N=3. A representative blot is shown. (C,D) U2OS cells were transfected with siRNA targeting MLH1 and MSH6 alongside a non-targeting control (siCTRL) prior to treatment with (C) IFNα (500 U/ml) or (D) IFNβ (100 U/ml) for 24 hours. Western blot analysis was performed to blot for PD-L1 as well as MSH6 and MLH1 to confirm knockdown. β actin was probed as a loading control. N=3. A representative blot is shown.

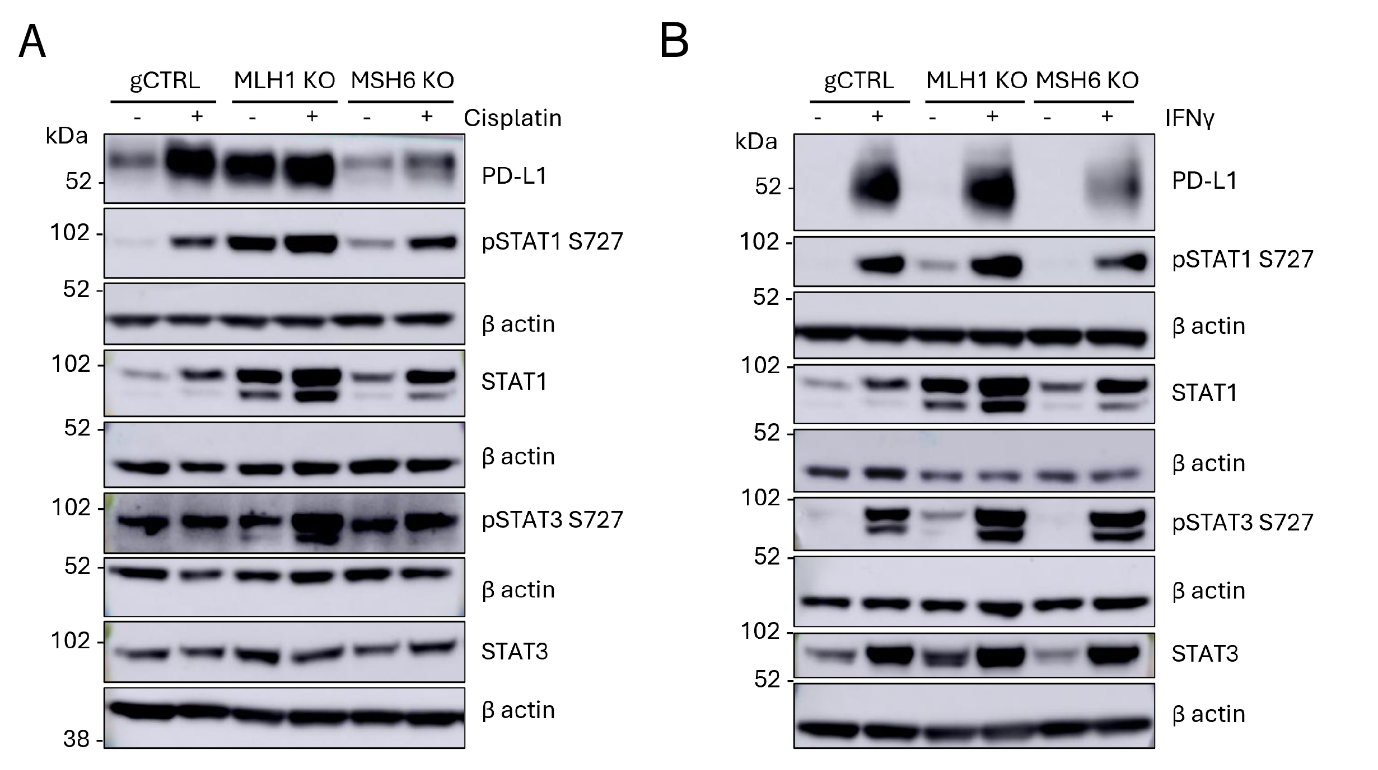

**Supplementary Figure 4. JAK-STAT signalling is not differentially regulated upon loss of MLH1 and MSH6.** CT26 gCTRL, MLH1 KO and MSH6 KO were treated with (A) cisplatin (2 μM) or (B) IFNγ (50 U/ml) for 48 hours. Western blot analysis was performed to probe for PD-L1 as well as STAT1/3 and their phosphorylated form (S727), which is required for the full transcriptional activity and biological function of STAT1/3. β actin was probed as a loading control. (A) N=3 or (B) N=2. A representative blot is shown.
